# Decision-making dynamics are predicted by arousal and uninstructed movements

**DOI:** 10.1101/2023.03.02.530651

**Authors:** Daniel Hulsey, Kevin Zumwalt, Luca Mazzucato, David A. McCormick, Santiago Jaramillo

## Abstract

During sensory-guided behavior, an animal’s decision-making dynamics unfold through sequences of distinct performance states, even while stimulus-reward contingencies remain static. Little is known about the factors that underlie these changes in task performance. We hypothesize that these decision-making dynamics can be predicted by externally observable measures, such as uninstructed movements and changes in arousal. Here, combining behavioral experiments in mice with computational modeling, we uncovered lawful relationships between transitions in strategic task performance states and an animal’s arousal and uninstructed movements. Using hidden Markov models applied to behavioral choices during sensory discrimination tasks, we found that animals fluctuate between minutes-long optimal, sub-optimal and disengaged performance states. Optimal state epochs were predicted by intermediate levels, and reduced variability, of pupil diameter, along with reduced variability in face movements and locomotion. Our results demonstrate that externally observable uninstructed behaviors can predict optimal performance states, and suggest mice regulate their arousal during optimal performance.

## Introduction

Behavioral and neural responses are notoriously variable across trials in both animals and humans (1). Recent studies have demonstrated that this variability is not random, but arises from changes in neural and physiological states (1-3). For example, arousal levels fluctuate on a moment-to-moment basis, and a significant fraction of neural and behavioral variability can be predicted by pupil diameter, which is tightly linked to arousal (4). Uninstructed movements also influence neural dynamics, as shown in the visual system (5-9) and reported ubiquitously across cerebral cortex (10-12). Methods for studying behavioral variations in relation to arousal vary but can lead to cohesive views of the nervous system and its function (13). For instance, the Yerkes-Dodson law, first proposed in 1908 and developed and modified through the years since, outlines an inverted-U relationship between arousal and performance on difficult tasks, with optimal performance occurring at intermediate arousal (14).

While some studies have successfully demonstrated a lawful inverted-U shaped relationship between behavioral performance and arousal, others have failed to achieve similar findings (4, 15). Uncovering the mechanisms of behavior is highly dependent upon detecting and understanding these behavioral and neural state-dependent processes. One possible reason for this inter-study variability is that certain sub-states of behavior were undetected or non-occurring. Further, these prior studies on the impact of arousal and movement consider moment-to-moment fluctuations, but do not address structured variations across time during task performance. Therefore, developing a robust strategy for detecting variations in behavioral strategy, and understanding how they interact, is an essential prerequisite to revealing the underlying neural mechanisms of behavior. A recent study (16) showed that mice and humans alike express a handful of discrete strategies during decision-making tasks, and switch between them within the same experimental session. This advances classical models of performance by accounting for structured fluctuations in engagement strategies. However, the precise relationship between arousal, movements, and discrete strategic performance states is not known.

Here, we address this fundamental question by linking transitions between decision-making strategies to changes in arousal (as measured by pupil diameter) and uninstructed movements (as measured by face video motion energy and locomotion speed). We identified epochs of optimal task performance and showed that trained animals maintain them for longer durations when compared to periods of sub-optimal strategic engagement. Consistent with the Yerkes-Dodson law, we found a striking inverted-U relationship between the likelihood of optimal state occupancy and pupil diameter and uninstructed movement in both auditory and visual discrimination tasks. Similarly, we found a U-relationship between the probability of task dis-engagement and pupil diameter/uninstructed movement. Furthermore, reduced variability in pupil diameter and movement measures signaled transitions into the optimal state and could be used to predict the onset and offset of decision-making states. Our results reveal that a significant fraction of behavioral variability can be accounted for through detection variations in sustained behavioral state/strategy, which can be predicted by shifts in a subject’s arousal and uninstructed movements, and further suggest that controlled arousal is key to maintaining optimal performance.

## Results

### Mice switch between several performance states during auditory and visual decision-making

To identify performance states/strategies explored by an animal during perceptual decision making, we trained mice for extended sessions on either an auditory or a visual stimulus discrimination task (Fig. 1A-C). Both versions of the task required mice to lick a left or right reward port to categorize the stimulus. For the auditory discrimination task, tone clouds were differentiated by the concentration of frequencies in high or low frequency bands, while in the visual task the angle of a drifting Gabor patch was categorized as closer to a vertical or horizontal orientation (Fig. 1B). We specifically sought to explore a broad range of arousal in our mice, from drowsy to highly aroused. Two factors of the task design were chosen specifically to promote this broad range of arousal during performance. First, mice were trained on a running wheel to allow high levels of arousal, such as those that occur with sustained running. Second, long inter-trial intervals (5 ± 2 seconds, Fig. 1C) were used to allow a reduction of arousal between stimuli, facilitating access to lower arousal levels, such as occurring during prolonged quiescence. Mice had three options available in each trial: lick left, lick right, or not respond. Indeed, we found that subjects sometimes did not respond following a stimulus, so we considered every stimulus presentation during analysis, including trials with no response (Fig. 1D).

**Fig. 1.**
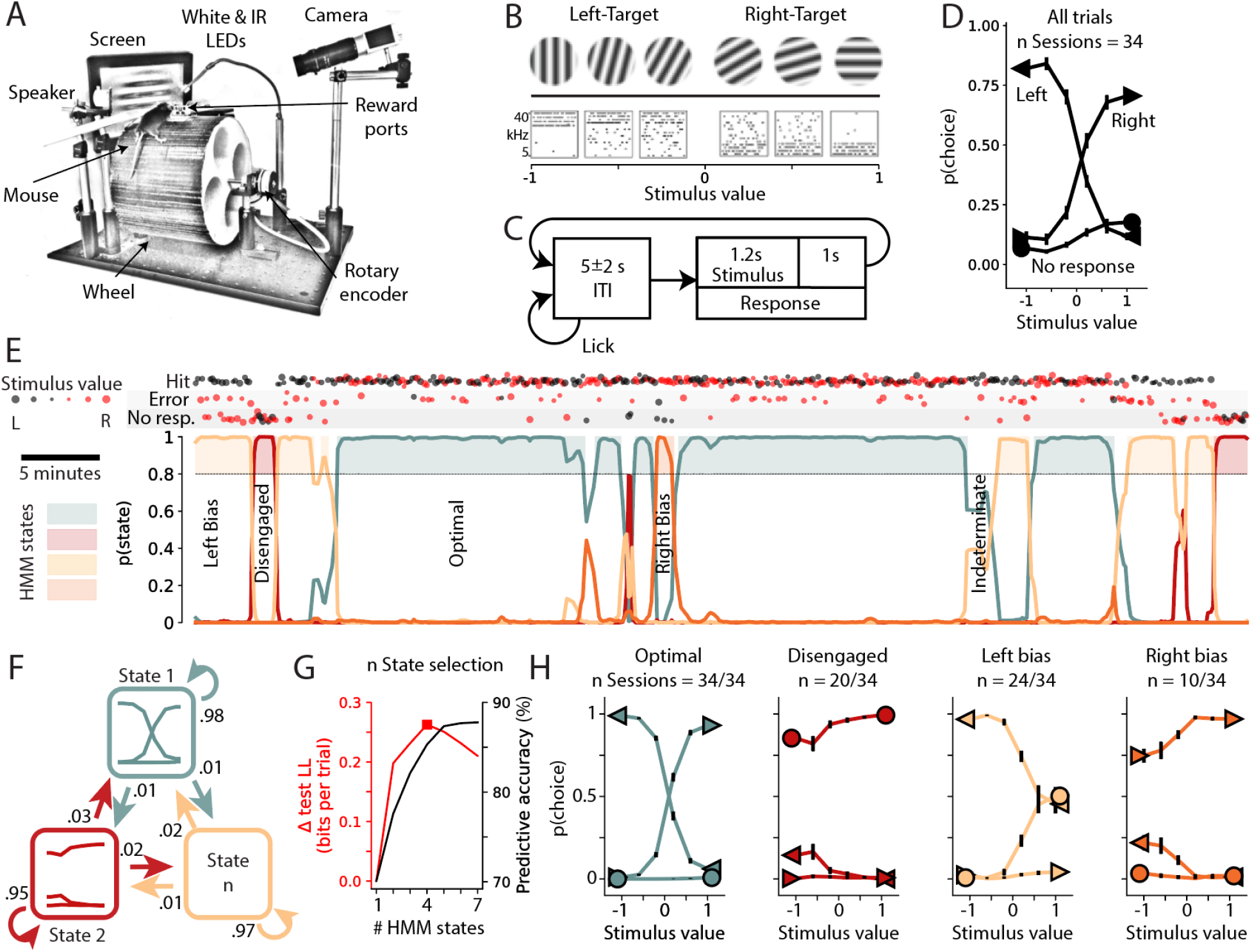
GLM-HMM reveals epochs of optimal performance amidst periods of sub-optimal strategies and complete disengagement. **(A)** During task performance mice were head-fixed above a running wheel in front of a screen and speaker. **(B)** Mice were trained to categorize the angle of a drifting Gabor patch as mostly vertical vs. horizontal, or the concentration of pure tones in a tone cloud as mostly high vs. low frequency. Negative stimulus values are associated with left-target stimuli and positive values with right-target stimuli. **(C)** Stimuli were presented after a random duration (5 ± 2s) of licking quiescence. Subsequently licking the corresponding (L/R) reward port resulted in water reward delivery. **(D)** Performance of an example mouse during the visual task, presented as the probability of each choice type (left (<), right (>), no response (o)) given each possible stimulus. **(E, top)** Structured fluctuations appear in trial outcomes during example session. Trial stimulus values are represented by marker size and color (size increases with ease of discrimination). **(E, bottom)** GLM-HMM state posterior probabilities capture fluctuations in performance dynamics. A confidence criterion of 80% was set for inclusion of a trial in a state for further analyses. **(F)** GLM-HMM, with generalized linear models of psychometric performance for each hidden Markov state, and state transition probabilities. **(G)** An appropriate number of states, denoted here by a red square, was determined using cross validated test-set log likelihood. Choice prediction accuracy continues to increase with number of states. **(H)** Performance of example mouse during trials sorted by GLM-HMM states. Some states occurred in only a subset of sessions (indicated by n sessions).

Examining the behavior during single sessions revealed that mice switch rapidly between epochs of varying stimulus-response contingencies (Fig. 1E, top). During the first five minutes the subject shown in this figure responded predominately to both left and right-target stimuli by licking to the left -thus exhibiting a strong left response bias. After several (30) trials of this left-bias behavior, the animal stopped responding to either left or right-target stimuli, and therefore was disengaged from the task (Fig. 1E, top). This 14-trial long period of disengagement was followed by another period of left-bias that lasted 29 trials. Suddenly, the animal started responding accurately to both left and right-target stimuli, in what we term as the “optimal state”. The subject remained in this state for a prolonged period of time (∼20 minutes). The remainder of this behavioral session could be roughly described as transitions between states characterized by either left-bias, right-bias, disengaged, optimal, or intermediate (indeterminant) states (Fig. 1E, top).

To more accurately and automatically capture these dynamics in performance, we used Hidden Markov Models with Generalized Linear Model emissions (GLM-HMM, Fig. 1F). Each HMM state (further referred to as a strategic or performance state) corresponds to a psychometric curve, captured by a GLM. GLM-HMM models have been previously used on two-alternative forced choice tasks with only two possible outcomes (L, R choices), leading to Bernoulli emissions (16). Here we extended this model to include multinomial emissions representing the three possible choices (L, R, no response). This proved crucial to capturing the full range of performance states, including disengagement. The model parameters include a transition probability matrix capturing the transition rates between different states, as well as GLM observation parameters representing the stimulus weights and bias of the psychometric curve in each state (Fig. 1F). The number of performance states for the model was determined for each mouse using cross-validation. For the example mouse shown in Figure 1G, this method yielded 4 discrete states. A final model was fit to all available data for a subject. The models yield posterior probabilities for each decision-making state in each trial, revealing long-lived states detected with high confidence and lasting for tens of consecutive trials (Fig. 1E, bottom).

Psychometrics generated from trials of each performance state yielded clearly interpretable and distinct task engagement profiles. For the representative subject presented in Figure 1, decision-making performance alternates between a state containing optimal performance, where responses were lawfully guided by the sensory stimulus, a disengaged state, with no response following stimuli, and sub-optimal left and right bias states, where the subject responded predominately in one direction regardless of the stimulus identity (Fig. 1H). From here on we refer to the state in which the animal is responsive and performance yields psychometric curves indicating responses are correctly guided by the stimulus, without significant leftward or rightward bias, as the “optimal state” (see Fig. 1H).

To compare the accuracy of this GLM-HMM technique to a model that assumes an animal does not switch between performance states, we use it to predict choices of the mouse on each trial. We again used cross-validation, and generated models trained on 80% of trials from each session, withholding blocks of 20% of trials as a test-set. Posterior probabilities of state occupancy for trials of test-set data were generated and used to predict the true choices based on the GLMs for each state. A one-state GLM-HMM, equivalent to assuming stationary performance during training, correctly predicted the choice of the mouse on 70% of trials, while the best-fit 4-state GLM-HMM correctly predicted 85% of choices (Fig. 1G).

To examine the generality and robustness of these observations, we fit GLM-HMM models separately to each of 13 mice that reached proficiency on the sensory discrimination task (8 auditory, and 5 visual task performers). While there was a diversity in the best number of states across subjects (3 to 5 states, Fig. S1), six stereotypical performance states emerged, as follows. Each subject exhibited both optimal (state 1) and disengaged (state 2) states, along with at least one additional sub-optimal performance state. In sub-optimal states, subjects either responded predominantly to the left (or right) regardless of stimulus value (yielding states 3 and 4 - left or right biased), or responded correctly to left-target (right-target) stimuli while withholding responses to right-target (left-target) stimuli (yielding states 5 and 6 - avoid right or avoid left, Fig. 2A). Subjects spent the largest proportion of trials in the optimal performance state (Fig. 2B; opt. vs dis. p = 2.4 × 10^−4^, opt. vs sub. p = × 10^−4^, opt. vs ind. p = 2.4 × 10^−4^, Wilcoxon signed-rank tests). Additionally, epochs of the optimal state lasted longer than those of sub-optimal states (Fig. 2C; opt. vs dis. p = 2.4 × 10^−4^, opt. vs sub. p = 2.4 × 10^−4^, Wilcoxon signed-rank tests). Finally, we determined the choice prediction accuracy for each mouse, and found the best-fit GLM-HMM correctly predicted 82.5 ± 4.3% of choices, a significant 17.7 ± 5.3% improvement over the classical static-performance assumption (Fig. 2D, p = 2.4 × 10^−4^, Wilcoxon signed-rank tests). These observations demonstrate that multinomial GLM-HMMs describe task performance better than classical static-performance models, that these models can be used to identify epochs of optimal performance. Using these models revealed that trained mice selectively maintain optimal strategic states longer than in sub-optimal strategies during task performance.

**Fig. 2.**
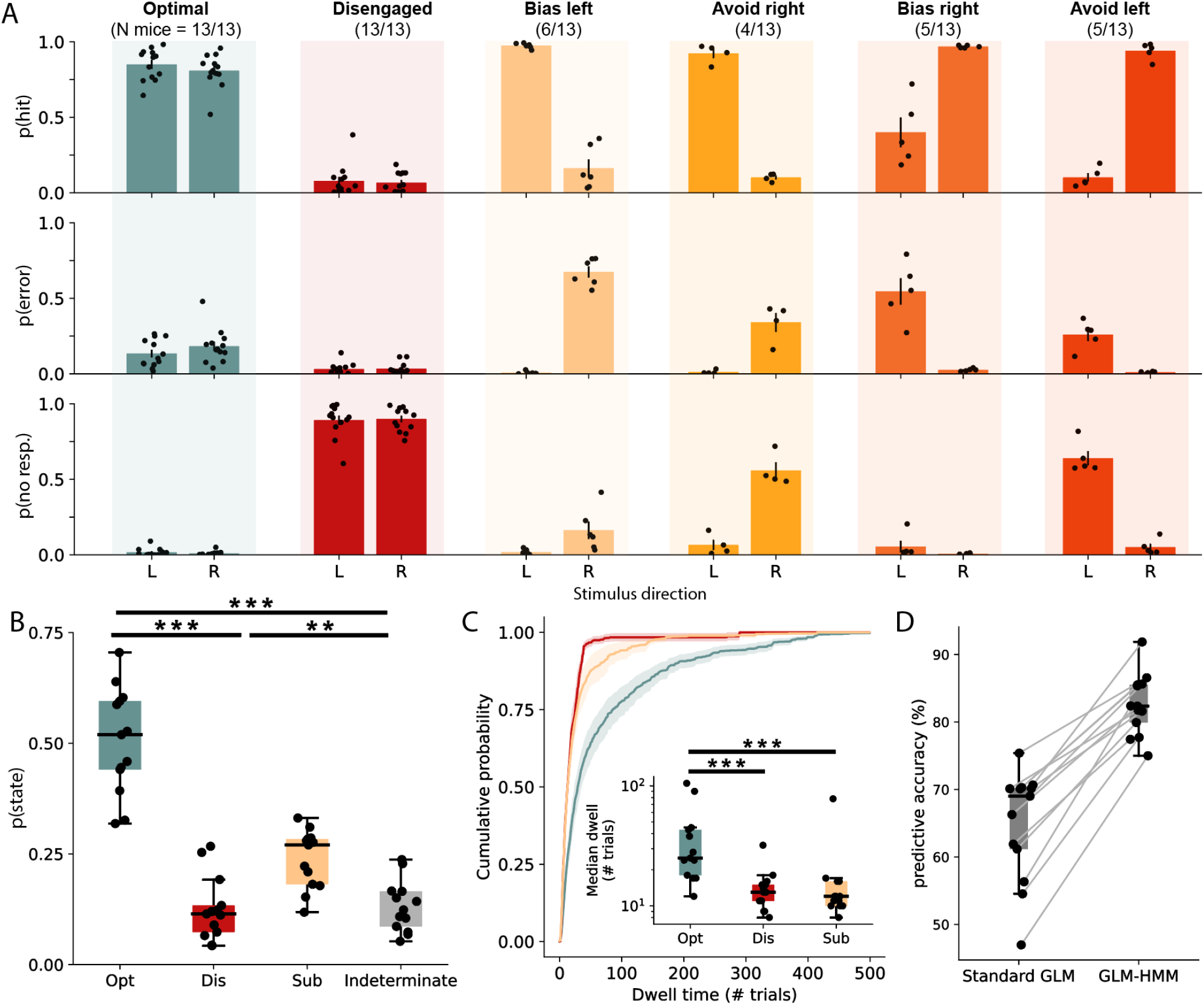
GLM-HMMs converge on six stereotypical performance states across mice. **(A)** Probability of hit, error, or no response for each mouse in each performance state. Each mouse had an optimal (L/R choice guided by stimulus) and disengaged (non-responsive to stimuli) state, along with one or more sub-optimal performance states. Sub-optimal states could be classified into four categories – bias left or right, where the mouse responds to the opposite stimulus with an error, and avoid left or right, where a stimulus of one direction was not responded to, while the other was responded to with a lick to the correct side. **(B)** Mice were in the optimal state around half of all trials. **(C)** Cumulative probability of performance state dwell times, averaged across individual mice. Inset: on each entry into an optimal state, mice remained in the optimal state for longer periods compared to other performance states. ** indicates p < 0.01, *** indicates p < 0.001. All data are represented as mean +/-SEM.

### Arousal and uninstructed movement are regulated during optimal performance

How do arousal and uninstructed movement relate to strategic states and their transitions? We hypothesized that there would be lawful relationships between arousal and movement (face, locomotion) measures, and discrete strategic performance states and their transitions. To examine this idea, we monitored measures of pupil diameter, face video motion energy, and locomotion speed of mice during task performance. Our analyses used the value of each measure immediately before the onset of a stimulus to determine their influence on variations in the subject’s response (Fig. 3A). A comparison between the time course of these arousal measure values and the HMM state dynamics within single sessions revealed precise relationships between these variables, providing support for our hypothesis (Fig. 3B). To quantify these relationships we examined how differences in pupil diameter and movements measures affected the probability of state occupancy (e.g., probability of being in the “optimal state”). Interestingly, we found a striking inverted-U relationship between probability of optimal state occupancy and pupil diameter, where the likelihood of being in the optimal state is highest at intermediate pupil diameters (Fig. 3C). The inverted-U relationship was found in nearly all subjects, and the pupil diameter that gave the highest probability for being in the optimal state was calculated for each (Fig. S2). Subjects performing the auditory task had significantly smaller optimal pupil sizes than those performing the visual task, suggesting optimal arousal levels may be task modality dependent (Fig. 3D; p = 0.013, Wilcoxon rank sum test).

**Fig. 3.**
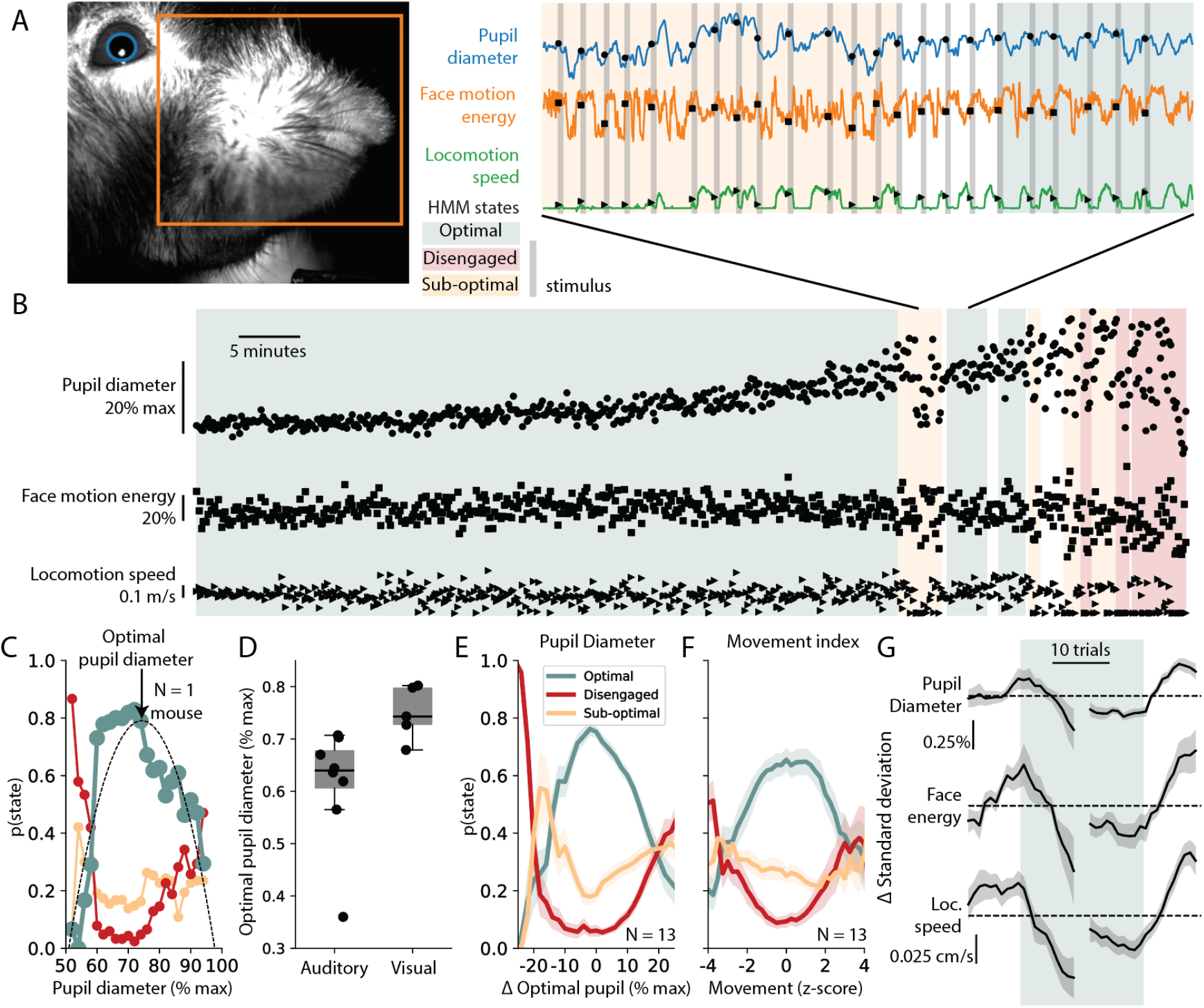
Optimal performance state is characterized by intermediate levels and lower variability of both pupil diameter and uninstructed movement. **(A)** Pupil diameter, face video motion energy, and locomotion speed before stimulus presentation are used for further analysis. **(B)** Example session shows shifts in arousal measures that coincide with GLM-HMM state changes. Note restricted variance and range of measures in optimal state as compared to disengaged or sub-optimal states. **(C**) Probability of GLM-HMM states across pupil diameters for an example mouse. Optimal state probability is fit with a polynomial function to estimate optimal pupil diameter for each mouse. **(D)** Optimal pupil diameter is significantly larger in mice performing the visual task when compared to those performing the auditory task. **(E)** Average probability of GLM-HMM states across all mice as a function of distance from optimal pupil reveals a robust inverted-U relationship with optimal state occupancy. **(F)** A movement index, combining face motion energy and locomotion speed, also has an inverted-U relationship with optimal state probability. There is a marked U-shaped relationship between both pupil diameter and movement index and the probability of being in the disengaged state. **(G)** Variability of all recorded measures decreases when entering the optimal state and increases when exiting this state. Plotted data represents the change in the standard deviation of each measure across the previous ten trials for trials surrounding optimal state transitions. Data were averaged across state transitions for each subject. Dashed line represents no change. Plots represent the mean +/-SEM across mice.

To visualize the relationship between pupil diameter and performance states across mice, we normalized the x-axis (pupil diameter) to the “difference from the optimal pupil diameter” for each subject to account for individual differences in the optimal pupil diameter (Fig. 3E). Similar to the inverted-U relationship between pupil diameter and probability of being in the optimal state, we found a prominent U relationship between the probability of being in the disengaged state and pupil diameter. At either low or high pupil diameters, there was a dramatic increase in the probability of mice being in the disengaged state, and a consequent large decrease in probability of the animal being in the optimal state (Fig. 3E). Individual analysis of movement measures showed that face motion energy had a largely monotonic relationship with probability of optimal state occupancy, and locomotion speed showed an inverted-U relationship across subjects (Fig. S3). The inverted-U relationship between the probability of optimal state occupancy and locomotion speed was less prominent than it was for pupil diameter, and was not present for all mice (Fig. S3).

Locomotion and facial movements are inter-related, but can convey independent information (e.g., while locomotion is always associated with facial movements such as whisking, facial movements often occur without overt locomotion). Owing to this, we combined these two measures into a single measure of uninstructed movement termed “movement index”. When face motion energy and locomotion speed were combined into a single movement index, a clear inverted-U pattern emerged, showing that optimal performance states occupancy peaked at intermediate levels of uninstructed movement (Fig. 3F; see materials and methods for movement index details). These correlations translate to a significant decrease in the distance from the optimal pupil diameter during optimal state occupancy as compared to sub-optimal or disengaged states, (Fig. S4; opt. vs dis. p = 2.4×10^−4^, opt. vs sub. p = 4.8×10^−4^, Wilcoxon signed-rank tests), a trend toward larger raw face video motion energy during optimal state occupancy (Fig. S4; opt. vs dis. p = 0.06, opt. vs sub. p = 0.04, Wilcoxon signed-rank tests), but no significant differences in raw locomotion speed (Fig. S4; opt. vs dis. p = 0.84, opt. vs sub. p = 1, Wilcoxon signed-rank tests).

We further hypothesized that performance states would differ not only in the values of arousal and movement measures, but also in their variability over the past ten trials. For all arousal measures recorded - pupil diameter, face motion energy, and locomotion speed - we found significantly reduced variability during optimal performance state occupancy as compared to sub-optimal or disengaged states (Fig. S4; pupil diameter: opt. vs dis. p = 2.4×10^−4^, opt. vs sub. p = 7.3×10^−4^; face motion energy: opt. vs dis. p = 3.4×10^−3^, opt. vs sub. p = 0.01; locomotion speed: opt. vs dis. p = 0.001, opt. vs sub. p = 0.048, Wilcoxon signed-rank tests). Strikingly, this decrease in variability predicts transitions into and out of the optimal state (Fig. 3B, G), and suggests arousal levels may be regulated during epochs of optimal task engagement.

### Arousal measures predict optimal performance state

To quantify the extent to which different performance states can be predicted based on the recorded arousal and movement measures, we implemented a cross-validated state classification analysis. For each subject, we trained a classifier to discriminate between optimal performance state and disengaged or sub-optimal states using the value and variability of each arousal measure as features. Figure 4A shows an example of such a classifier using the value and trial-to-trial variability of pupil diameter. We found that for all subjects, performance states could be significantly predicted using the recorded arousal measures with high accuracy (opt. vs dis. = 88.2% ± 6.1; opt. vs sub. = 77.7% ± 8.6; Fig. 4B-C; significance was assessed by comparing empirical classification accuracy to surrogate datasets obtained by shuffling class labels in the training set). To assess the contributions of individual arousal measures to performance state decoding, subsets of measures were shuffled in the training sets of the classifiers. Pupil diameter (value and trial-to-trial variability) was the best individual raw measure for state classification (Fig. S5; pupil vs face motion energy p = 3.9×10^−3^, pupil vs locomotion p = 0.106, Wilcoxon signed-rank tests). Interestingly, we found that the best predictor of performance state was the computed movement index incorporating face motion energy and locomotion speed, which encoded more information about states compared to pupil diameter (Fig. 4C; pupil vs movement index p = 0.024, Wilcoxon signed-rank tests).

**Fig. 4.**
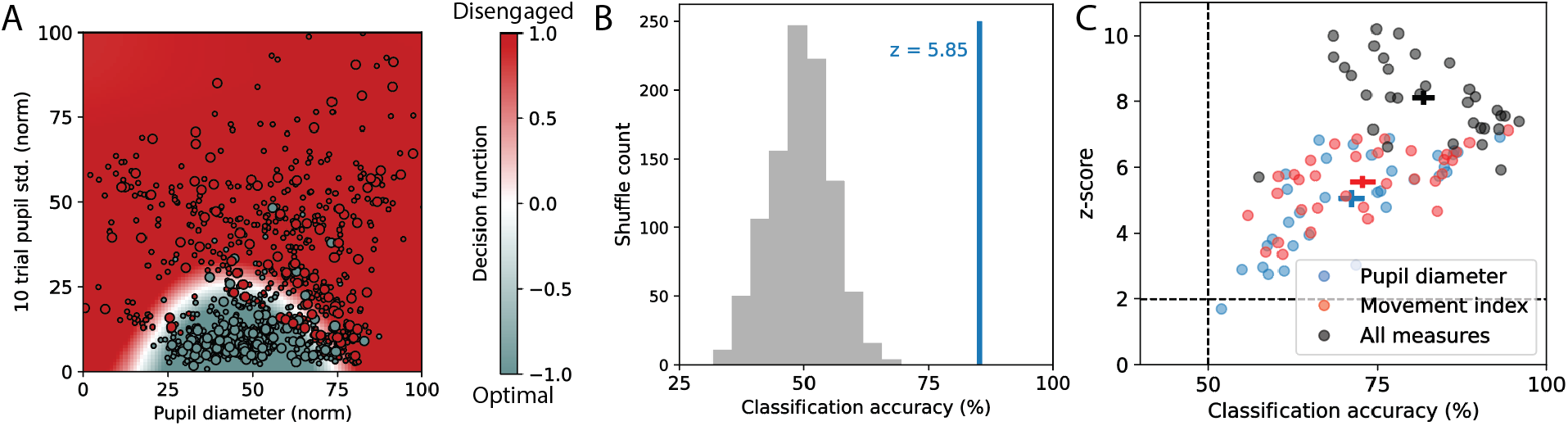
Arousal and movement measures effectively predict optimal state occupancy. **(A)** Example decision function of a state vector machine (SVM) classifier with a radial basis function kernel using pupil measures of diameter and trial-to-trial variability to predict performance states. One of five cross-validated folds shown. Color bar represents decision function confidence, with white representing the decision boundary. Small circles: training set, large circles: test set. **(B)** Example of z-scoring test set accuracy of performance state decoder trained on empirical data (blue line; one fold from panel A) against class-shuffled data (gray distribution). **(C)** Performance state classification accuracy and z-scores for each classifier. To determine the accuracy of classification using the value and variability of individual measures all other measures were shuffled. All data are represented as mean +/-SEM.

## Discussion

In this study we provide evidence that a significant fraction of behavioral variability arises from rapid transitions between identifiable substates, that pupil diameter and uninstructed movements can accurately predict these substates and their transitions (including epochs of optimal performance), and that controlled arousal is key to maintaining optimal performance. First, using GLM-HMM modeling of task performance we show that well-trained mice alternate between discrete strategic performance states, and accounting for these states significantly improves the accuracy of behavioral modeling. Our methods, including trials where the subject does not respond, allowed us to go beyond previous reports^16^ and identify disengaged and other low performance states in addition to optimal performance states. Interestingly, we found that mice selectively maintained optimal states for longer durations than sub-optimal and disengaged states (Fig. 1D, 2C). Second, consistent with the classical Yerkes-Dodson law, we demonstrate that optimal task performance occurs at intermediate arousal levels during sensory discrimination tasks (Fig. 3E, S2). While previous reports demonstrated this relationship during auditory detection (4), it has not been reported during visual tasks (15). Here we find an inverted-U relationship during both auditory and visual discrimination task performance. There was a significant difference in optimal pupil diameters between sensory modalities, suggesting modality specific modulation of arousal during optimal performance. Third, we found significantly lower trial-to-trial variability in both pupil diameter and movement measures during optimal performance, with rapid shifts in variability at transitions into and out of the optimal state (Fig. 3G, S4). Finally, we found that arousal measures, taken prior to stimulus presentation, can be used to accurately predict the discrete task performance states produced by GLM-HMM modeling (Fig. 4C). By using GLM-HMMs, we extend analysis of arousal measures beyond average characterizations of a moment-to-moment performance-arousal relationship in mice and suggest strategic regulation of arousal levels during optimal performance.

Hidden Markov models are a crucial tool for inferring performance states. Here we used recently implemented GLM-HMM models (16, 17) and advanced their utility to account for stimuli not responded to by the subject. This extended model accounts for additional behavioral states present during decision making in mice, in particular revealing states of disengagement and selective avoidance of a particular stimulus category (*i*.*e*., left-or right-target). Using an individual model for each subject accounted for differences in individual psychometric response patterns and state transition probabilities. Despite individual differences, when comparing across subjects a consistent view of task performance emerged, with a limited number of easily classifiable, congruent strategies in the population. In the current study, we observed arousal related differences between optimal and sub-optimal strategic states, but different performance states may additionally have markedly different neural activity and/or functional connectivity correlates. Studying neural data regarding decision making could benefit from comparing not only hits and errors, but also comparing differences between similar trial outcomes during various strategic performance states.

Arousal and movement can be indexed by numerous, often correlated measures (1). Here we use simple and accessible measures of pupil diameter, face video motion energy, and locomotion speed. Head-fixed mice on a running wheel can exhibit a broad range of arousal and movement, from sleep to rapid locomotion. To promote a broad range of arousal in our mice, we both allowed mice to locomote rapidly on a running wheel during task performance (therefore allowing access to higher levels of arousal and uninstructed movement) and used long inter-trial-intervals, to promote periods of behavioral quiescence and task disengagement. Preliminary results indicated that short inter-trial intervals typically resulted in a lack of lower levels of arousal (*e*.*g*., smaller pupil diameter during behavioral quiescence) and biased our results towards intermediate to high arousal levels. These factors were crucial for observations of an inverted-U relationship between optimal state probability and pupil diameter (Fig. 3E). We additionally saw an inverted-U relationship between a movement index combining face motion energy and locomotion speed with optimal state probability, similar to that of pupil diameter (Fig. 3F). The measurement of face motion energy individually showed a monotonically increasing relationship with optimal performance state occupancy. Onset of facial movements tracks rapid shifts in intracellular (6, 18, 19) and neuromodulatory neural dynamics (19-21) but lacks fidelity during prolonged bouts of movement and locomotion, when face movement is ubiquitous. Conversely, locomotion speed does not capture the onset of whisking and movement, but provides a continuous measure of high-energy movements. Thus, our facial motion energy measure is well-suited for detecting both lower levels of uninstructed movements (*e*.*g*., movements that do not include the large skeletal muscles of the limbs and body), while locomotion speed is better suited for detecting higher levels of these movements. While some mice had an inverted-U relationship between these individual measures and optimal performance state, combining the two movement measures allowed for one index to account for a wide range of movement levels, and resulted in an inverted-U relationship with optimal performance state across all mice. Based on these factors we consider pupil diameter to be the best easily accessible, individual measure of arousal. While complex analysis of video data could lead to more precise measures of unique movements profiles for individual subjects basis, the simple movement index used here is sufficient to make accurate predictions of task performance state and corroborates a long standing physiological phenomenon first described by Yerkes and Dodson (14).

What factors contribute to the inverted-U phenomenon during task performance? In the auditory cortex, membrane potential dynamics at intermediate arousal levels are ideal for stimulus detection (4). Task engagement also influences auditory processing in multiple brain regions, and changes are congruent and often overlapping with pupil-linked effects (22). In this case maintaining an intermediate arousal level could directly improve discriminability of neural representations and task performance. However, in the visual system, locomotion increases neural responses along with stimulus decoding accuracy using neural activity at the population level (5, 23-25). How this relates to behavioral performance is not clear, as rodent performance on visual tasks generally lags behind population level neural decoding potentials in the visual system of mice (26). While individual neural responses in the visual system vary in their relationship to locomotion speed (27, 28), decoding accuracy has often only been tested with a binary classification of states (stationary vs locomoting). Binary classification of locomotion in some instances indicates locomotion is beneficial for task performance, while others have found it to be detrimental (15, 29). In the present study, the probability of both task disengagement and sub-optimal strategic states increased at higher levels of both the continuous pupil diameter and movement indexed, leading to decreased performance at high arousal/movement levels across both auditory and visual modalities. Crucial to this observation is that movement is measured on a continuous scale. In addition, effects on stimulus decoding accuracy have only been performed in passive contexts and are restricted to the visual cortex. While arousal-dependent cortical decoding capacities may contribute to behavioral performance, other brain regions receiving their signals may have different arousal-dependent dynamics, where heightened arousal may inhibit optimal performance.

Regulation of arousal to maintain ideal levels for decision making may lead to increased performance. During difficult tasks, external feedback based on optimal neural activity in humans can increase performance and decrease arousal as measured by decreases in pupil diameter and increases in heart rate variability (30), consistent with the right half of the Yerkes-Dodson relationship. Internal factors may also regulate arousal. Development during childhood and adolescence leads to self-regulation of arousal, contributing to executive function in humans (31-33). This developmental phenomenon is not unique to humans -in mice projections from the prefrontal cortex (PFC) to the serotonergic dorsal raphe nucleus (DRN) increase through adolescence, and their emergence coincide with increases in persistence during active foraging (34). Recent work has shown humans can gain volitional control of pupil size through training, systematically regulating neural structures related to arousal (35). In the present study, in addition to finding an ideal range of pupil diameters for optimal task performance, we report a decrease in trial-to-trial variability of all recorded arousal measures during epochs of optimal performance states, and differing optimal pupil diameters based on sensory modality (Fig.s S4, 3D). While we do not present direct evidence of volitional control of arousal in mice, the arousal regulation shown during optimal engagement states opens questions regarding the mechanisms that contribute to this effect. While strategic control of movements may account for much of the arousal changes, further study is warranted into the use and development of regulatory mechanisms and contextual control of arousal.

What is the neural mechanism underlying performance state switching and its arousal-induced regulation? Fluctuations in pupil diameter are correlated with activity in arousal-linked neuromodulatory centers in mice, non-human primates and humans (20, 36, 37). The PFC is well situated to be an orchestrator of neuromodulatory centers linked to arousal and pupil size. In mice, PFC projections target both serotonergic and GABAergic populations in the DRN, and electrical stimulation of the PFC modulates DRN activity (38-40). Direct activation of serotonergic cells in the DRN sustains arousal, slowing pupil constriction (41). Similarly, direct stimulation of the noradrenergic locus coeruleus (LC) leads to increases in pupil diameter, and stimulation of PFC projections to inhibitory populations surrounding the LC leads to pupil constriction (42). PFC projections can act as dynamic regulators for key arousal-linked neuromodulatory centers, and their coordination could maintain optimal arousal levels during task engagement. Further work is necessary to determine the activity and influence of such projections during various task engagement states.

## Materials and Methods

### Animals

All procedures were carried out with approval from the University of Oregon Institutional Animal Care and Use Committee. Animals (Female and male mice, 8-15 weeks at time of surgery) were of C57BL/6J background purchased from Jackson Laboratory and bred in-house, including wild-type C57BL/6J, transgenic Cre and tTA driver lines (CaMK2-Cre, Jax #005369; CaMK2-tTA, Jax #007004), and fluorescent reporter lines (tetO-GCaMP6, Jax #024724; TIGRE2.0 GCaMP6s Jax #31562). The mice were kept on a reverse light cycle and had *ad-libitum* access to food and water until time of behavioral training.

### Headplate implantation

All surgical procedures were performed in an aseptic environment with mice under 1-2% isoflurane anesthesia with an oxygen flow rate of 1.5 L/min, and homeothermic maintenance at 36.5° C. Mice were administered systemic analgesia (Meloxicam SR: 6 mg/kg, Buprenorphine SR: 0.5 mg/kg) and a fluid supplement (1 ml lactated ringer’s solution) subcutaneously. Fur was removed across the dorsal and right temporal surfaces of the skull, and the skin was sterilized with povidone/iodine solution followed by isopropyl alcohol three times over. The skin, connective tissue, and part of the right temporalis muscle were removed, and the exposed skull was cleaned. A custom-designed headplate was affixed to the skull using dental cement (RelyX Unicem Aplicap, 3M), and skin was affixed to the outside edge of the headpost as necessary (Vetbond, 3M). The exposed skull was covered using cyanoacrylate (Slo-zap, Zap), and protected with a silicone elastomer (Kwik-Sil, World Precision Instruments). Mice recovered for three days in an incubator recovery chamber, and lactated ringer’s solution was administered as necessary.

### Task details

Data collection and stimulus presentation was conducted using custom LabView (National Instruments) scripts. Mice were headfixed on a running wheel and trained on either an auditory or a visual stimulus discrimination task to receive water rewards. Mice made a binary classification of stimuli, and reported their choice by licking one of two reward ports (left and right) spaced 500 µm apart.

For the auditory task, stimuli consisted of tone clouds with three concurrent streams of tones, where each tone lasted 30 ms and had a frequency selected between 5 – 40 kHz. Tone clouds were differentiated by selecting a varied proportion of tone frequencies in either the bottom (5-10 kHz) or top (20-40 kHz) octave of the range. The remaining tones were randomly distributed across the rest of the frequency range. Auditory stimulus values are defined based on the proportion of tones in the right-target octave (e.g. high frequency) minus the proportion of tones in the left-target octave (e.g. low frequency), such that -1 represents a tone cloud with all tones in the left-target octave, 1 represents a tone cloud with all right-target octave tones, and 0 represents a stimulus with an equal number of tones in each target octave. Tones were calibrated to 60 dB SPL and waveforms were generated (NI PXI-4461, National Instruments) at 200 kHz sampling rate, conditioned (ED1, Tucker Davis Technologies), and transduced by electrostatic speakers (ES1, Tucker Davis Technologies).

For the visual task, stimuli consisted of drifting Gabor patches displayed on an LED screen with a refresh rate of 30 Hz. Stimuli had a constant mean luminance, matched to the static gray background displayed between stimuli. Each Gabor had 0.08 cycles per degree of visual field and drifted at 1.5 cycles per second. Gabor were differentiated by being closer to a horizontal (e.g., 0, 18, or 36 degrees) or vertical (e.g., 90, 72, or 54 degrees) orientation. Visual stimulus values were defined between 0 and 1 by the normalized difference of their angle from 45 degrees, and signed in accordance with their directional representation (negative if left, positive if right).

Stimuli (high/low tone cloud or horizontal/vertical gabor) were randomly assigned to left or right identities for each subject at the beginning of training. In both tasks stimuli lasted 1.2 seconds and were presented with an inter trial interval (ITI) of 5 ± 2 seconds. Licking of reward ports was electrically monitored at 1 kHz (USB-6008, National Instruments). If mice responded to stimuli by licking the appropriate reward port during the stimulus or within 1 second of the stimulus offset, the trial was considered a hit and a 3 μL reward was delivered through the port by a syringe pump (NE-500, Pump Systems Inc.). If the incorrect port was licked first during the stimulus or response period the trial was considered an error. If no response was made the trial was considered to have no response.

### Training & inclusion criteria

After postoperative recovery, mice were weighed for three days to establish a baseline weight before beginning a water regulation protocol. After 3-5 days, mice had reached a steady weight on water regulation and began behavioral training. Mice were habituated to human handling, head fixation, locomotion on the wheel and collecting water from the reward ports. Reward port position was adjusted by a manual xyz manipulator. To ensure consistent lick spout positioning, a 3D printed jig was made for each mouse and used to align reward ports at the beginning of each training session.

Behavioral training was conducted using the following stages and progression criteria:

Stage 1 - To encourage licking and associate left and right ports with the appropriate directional stimuli, the left or right port was randomly ‘armed,’ so a lick would prompt appropriate (value -1 if left, 1 if right) stimulus presentation, and water reward delivery. The stimulus and reward were spontaneously delivered after 20 seconds if the subject did not lick the armed port. The ITI was set to 2 seconds to promote consistent licking. After 75 such trials in a session, stimuli were presented without a reward, and a reward was delivered if the subject licked the correct port within the response period. If subjects achieved 1500 total licks to each port, and >150 hits in a session, they progressed to stage 2 in their next training session.

Stage 2 - To promote licking only following stimulus presentation, the ITI was extended to 5 ± 2 seconds, and was reset following aberrant licking for all following stages. Only the easiest stimulus values were used (−1 or 1), and a reward was delivered following any lick to the correct port during the response period. If subjects failed to respond to 10 consecutive trials, a free reward was delivered with the stimulus. If subjects achieved >50% hit rate on both ports, with no less than 30% of licks to each port and >150 hits, they progressed to stage 3.

Stage 3 - To refine choice behavior, a reward was only delivered if the first lick following stimulus presentation was to the correct reward port. To prevent mice from only responding at one reward port, probabilistic bias correction was added, so that more target stimuli of one direction would be presented if that port was being neglected. If subjects achieved >60% hit rate on both ports and >150 hits, with no less than 33% of hits at each port they progressed to stage 4.

Stage 4 - More difficult stimuli were introduced, and probabilistic bias correction was reduced. If subjects achieved >60% hit rate on both ports and >200 hits, with no less than 33% of hits at each port, they progressed to stage 5.

Stage 5 - Bias correction was removed and three difficulties of stimuli are presented per direction. A subset of mice were advanced to stage 6 after at least 10 consecutive stage 5 training sessions.

Stage 6 - Half of the stimuli were of the un-trained sensory modality, presented as distractors. There was no discernible change in task performance, and response rates to the distractors were <10%.

If subjects failed to meet the advancement criteria of a lower training stage for consecutive sessions, they were returned to that training stage. For further analysis we included sessions from stages 5 and 6 with at least 100 rewards delivered, balanced such that no less than 20% of rewards were delivered in each of the left and right ports. 13 of 31 mice had 10 or more sessions that met inclusion criteria, with a total of 381 of their 500 stage 5 and 6 training sessions included for further analysis. Mice that reached proficiency took 21 ± 8 training sessions to reach stage 5.

### Modeling task performance

To test the hypothesis that animals switch between discrete decision-making states within single sessions, we developed a hidden Markov model with multinomial Generalized Linear Model observations (GLM-HMM). The multinomial GLM observation, parameterized as

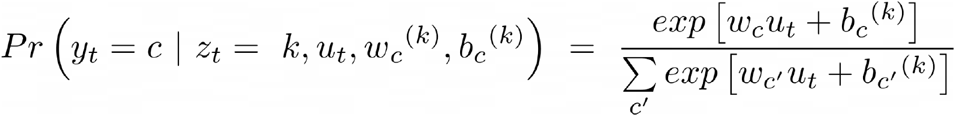

is the set of three psychometric curves representing the probability of choosing actions c=Left, Right, No response in each trial t, given stimulus *u*_*t*_ and the hidden state *z*_*t*_ = *k*. Each hidden state represents a different performance state, including one optimal, and several sub-optimal or disengaged states. A model with K hidden states is described by the following parameters: a KxK transition probability matrix *A*_*kk*_, representing the probability of switching between different states at each trial *t*; a K-dimensional vector representing the initial state probabilities *π*^(*k*)^; and the observation parameters comprising weights *w*_*c*_^(*k*)^ and biases *b*_*c*_^(*k*)^ for each of three multinomial categories *c=L,R,Nr*, the latter corresponding to three choices Left, Right, and No response, available to the subject in each trial. We fit a multinomial GLM-HMM to trials from individual subjects using the Expectation-Maximization (EM) algorithm to maximize the log-posterior and obtain the optimized parameters *θ* = {*w*_*c*_^(*k*)^, *b*_*c*_^(*k*)^, *A*_*kk*_, *π*^(*k*)^}. Model selection for the number of states was performed using 5-fold cross-validation across sessions of an individual subject. A model was fit to the concatenated trials of the training set, and the log-posterior of the test set was estimated (normalized by the number of trials per test set). Because the EM may lead to local maxima of the log-posterior, for each choice of number of states, the EM algorithm was performed 10 times starting from random initial conditions. We performed model selection in two alternative ways: either by maximum likelihood estimation (MLE); or by maximum a posteriori (MAP, including a Gaussian prior on the weights with mean zero and variance equals to 2, and Dirichlet prior on transition probabilities with alpha=2, see ref 16 for details on the procedure). The best number of states was chosen as the maximum of the plateau of the test MLE or test MAP log likelihoods. We then fit a single model to the time series of the observations and inputs concatenating all sessions from a subject, optimizing the model parameters *θ* using MLE.

For all further analysis we set an 80% state probability criterion for inclusion of a trial within a performance state and consider all other trials as in an indeterminate state. Performance states from the final models were distinct and clearly interpretable, and classified as one of six stereotypic states: optimal (responses congruent with stimulus identity), left bias (leftward licks regardless of stimulus), avoid right (left lick to left-target, no response to right-target stimuli), right bias (rightward licks regardless of stimulus), avoid left (right lick to right-target, no response to left-target stimuli), or disengaged (no response to any stimuli).

### Recording Arousal Measures

All data collection was conducted using custom LabView scripts. While headfixed, subjects were free to locomote on top of a cylindrical wheel with a 15 cm diameter and 20 cm width. The axle of the wheel was connected to a rotary encoder (Model 15T/H, Encoder Products Company), which was used to calculate locomotion speed.

A CMOS camera (Teledyne G3-GM11-M2020, or Basler ace acA780-75gm) with an affixed lens (TEC-M55MPW, or Navitar NMV-50M23) and IR filter (MIDOPT BN810-46, or Thorlabs FGL780) was pointed at the face of the subject. The face was illuminated with an infrared LED (Digi-Key TSHG8200, 830 nm) adjusted to provide even illumination of the face and eye, and with a white LED (RadioShack 5 mm 276-0017), adjusted so the pupil would have large dynamic range, while not being obscured by the eyelid when maximally dilated. Images of the face were acquired at 30 Hz throughout task performance, and time stamps for each frame were saved at time of acquisition.

Online pupil diameter and face motion energy estimates were made using LabView software, and post-hoc analysis was done using custom python scripts as necessary. Face motion energy was calculated within a rectangular ROI anterior to the eye (Fig. 3A). The absolute value of the frame-to-frame change in pixel intensity was averaged within the ROI and normalized to the maximum within a session. To calculate pupil diameter an ROI around the eye was selected and the image was binarized with a manually selected threshold. The contour of the pupil was extracted and the long axis of a fit ellipse was recorded as the pupil diameter. Pupil diameters were normalized to the maximum within a training session. Pupil diameters were manually verified, and if accurate measurements could not be acquired, frames were dropped and values interpolated if gaps between data points were less than 200 ms.

All measures were smoothed with second order Savitzky Golay filters with 500 ms (face energy and locomotion speed) or 1 second (pupil diameter) windows and upsampled to 1 kHz. In order to determine the influence of arousal and movement on performance, measure values concurrent with the beginning of stimulus presentation were used for further analysis. In addition to the raw values, the standard deviation and coefficient of variation of the measures’ values over the past 10 trials were calculated for each trial.

The measures of face motion energy and locomotion speed individually account for qualitatively distinct magnitudes of movement in mice during head fixation. Face motion energy effectively captures change in low-medium ranges of movement while the subject is stationary (not walking), and locomotion speed effectively differentiates medium-high output movements. In order to create a single value representative of a broad range of movement, the values of face motion energy and locomotion speed were z-scored and summed to create a single continuous movement index. The same was done for the past 10 trial standard deviation values.

### Determining optimal pupil diameters

Probability of a performance state was calculated in relation to each arousal measure by binning trials by measure values and determining the proportion of trials in each performance state. In order to determine the optimal pupil diameter for each subject, polynomial functions were fit to the optimal state occupancy probability across pupil diameters. Polynomial degree for a fit was selected by determining the elbow of the test set r^2^ increase function with 5-fold cross validation. A final polynomial was fit to all data, and the optimal pupil diameter was determined for each subject as the maximum of the fit curve within the range of the true data. The unique optimal pupil value for each subject was used to align pupil values for visualization, and to calculate the absolute difference from the optimal pupil diameter for each trial.

### Performance state classification from arousal measures

We used cross-validated classifiers (support vector machines with a radial basis function kernel) to discriminate trials from the optimal performance states against trials of each of the disengaged and sub-optimal states (binary classifications) using six features: the 3 recorded behavioral measures and their variability over the 10-past-trials. Classification was performed on a randomly selected equal number of trials of each performance state using 5-fold cross-validation. Accuracy is reported as the mean accuracy of test set classification across five such folds. To test the significance of the reported accuracies, we z-scored the empirical accuracy against 1000 classifiers trained on surrogate data obtained by randomly permuting class labels (43). To determine the contribution of features to classification accuracy, we compared the performance of the classifiers trained on the empirical data to those trained on surrogate data obtained by shuffling the labels for a subset of features across trials within the training set.

### Statistical Analysis

Sample sizes were not predetermined with statistical methods but are similar to other studies in the field. Data are reported as the mean ± standard deviation in the text, and are plotted with standard error of the mean. Two-tailed Wilcoxon signed rank (paired) or rank sum (unpaired) statistical tests were used to avoid assumptions regarding the normality of data distribution. Individual data points are shown when possible.

## Acknowledgments

We thank Benjamin Switzman & Austen Chesbro for assistance with behavior training, and Dennis Nestvogel, Lindsay Collins & Evan Vickers for fruitful discussions regarding this work and feedback on the manuscript. We further thank Evan Vickers for sharing the headplate design and training for surgical implants.

## Author contributions

Conceptualization: DH, LM, DAM, S.J

Methodology: DH, LM, SJ

Software: DH, LM, SJ

Formal Analysis: DH

Investigation: DH, KZ

Writing - Original Draft: DH, LM

Writing - Review & Editing: DH, LM, DAM, SJ

Visualization: DH

Supervision: LM, DAM, SJ

Funding Acquisition: LM, DAM, SJ

## Data and materials availability

All data needed to evaluate the conclusions in the paper are present in the paper and/or the Supplementary Materials. Analysis code is available at https://github.com/sjara/uobrainflex/tree/master/hulsey2023 and data will be shared upon request.

## Supplemental Figures

**Fig. S1.**
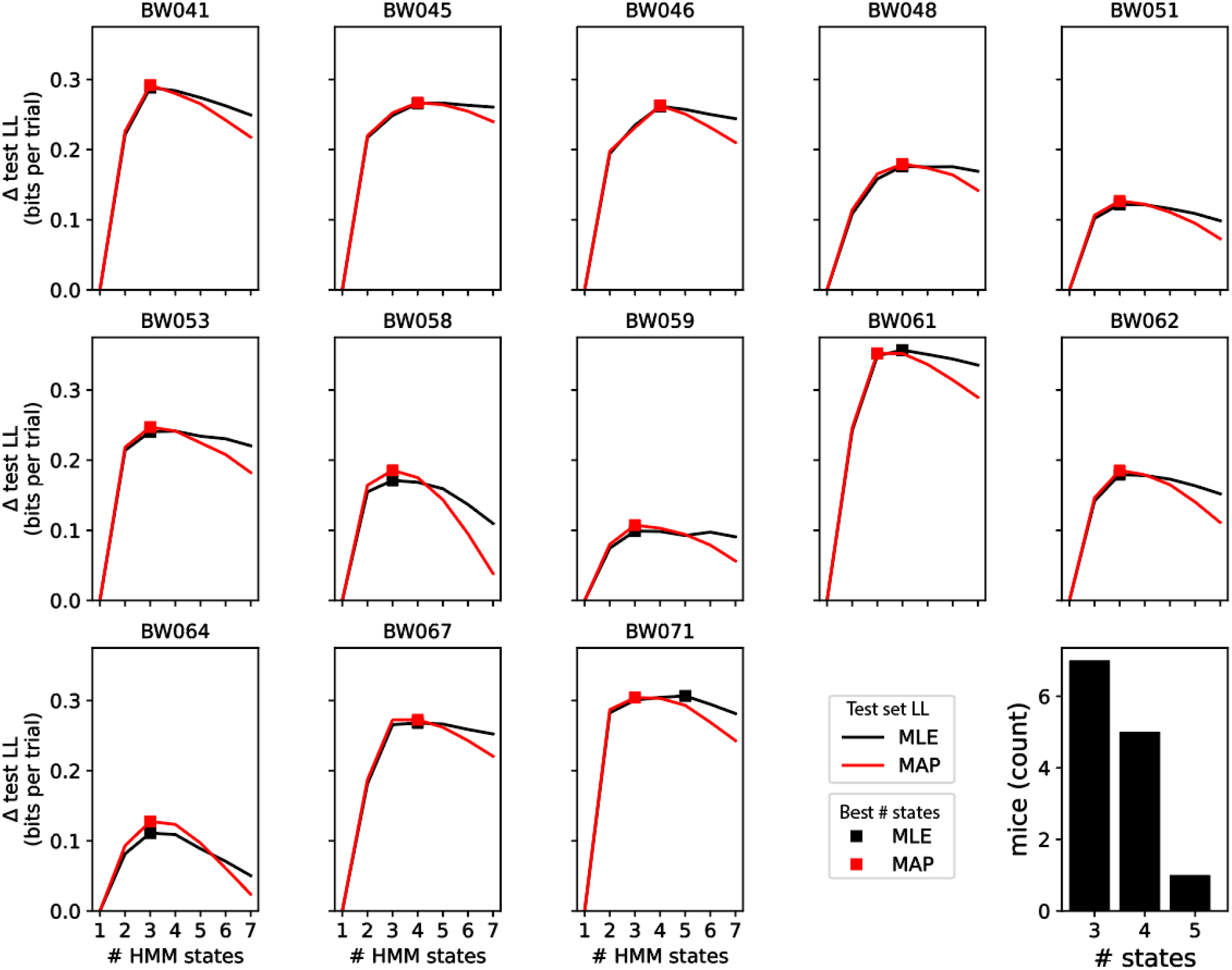
Selection of number of HMM states. State selection was performed using five-fold cross validation of test set log likelihood (LL), training models on 80% of sessions, and testing the fit on the remaining 20%. Both MLE and MAP methods were used (see materials and methods), with ten initializations per state and model type, and the maximum LL acquired was kept. The plateau of the average LL across the five folds was considered the appropriate number of states to use for final model fitting. In case of discrepancies between MLE and MAP states, the larger number of states was used.

**Fig. S2.**
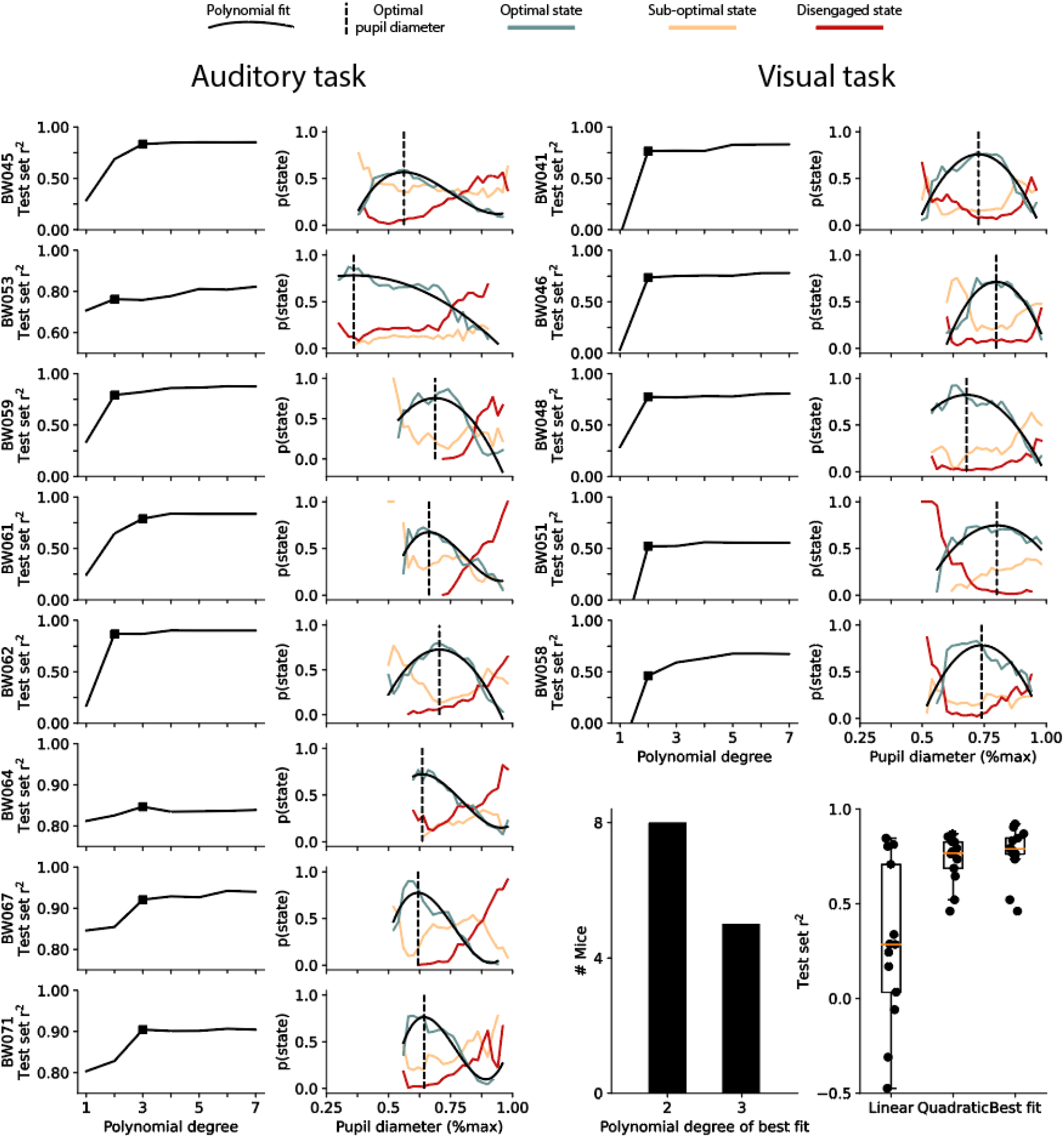
Optimal pupil diameter can be calculated for individual subjects. Polynomial functions were fit to the optimal state occupancy probability data for each individual mouse. Polynomial degree was selected using 5 fold cross-validation, and the elbow of the function was used to fit a final curve to all data. For the majority of mice, a quadratic function was sufficient to explain most of the variability. A cubic function was better fit in individuals with long tails to the inverted-U relationship. The optimal pupil diameter for each subject was defined as the maximum along the fit curve within the range of the true pupil data.

**Fig. S3.**
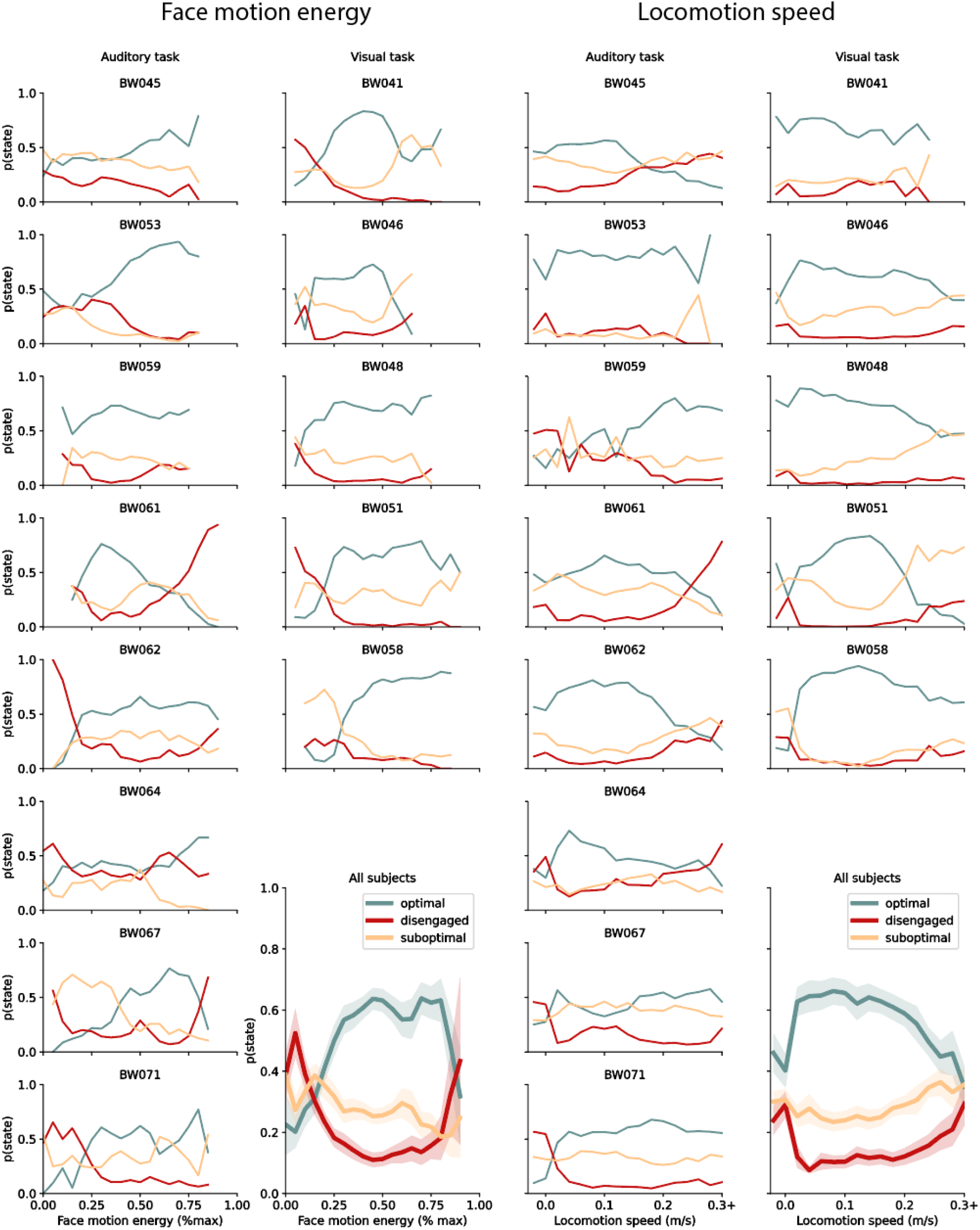
Relationship between optimal state occupancy and individual movement measures are variable. Individual mice have inverted-U relationships between face motion energy and locomotion speed. However, the relationship is inconsistent, and not found in all subjects. Across subjects, face motion energy forms a largely monotonic relationship, while locomotion speed has a moderate inverted-U relationship.

**Fig. S4.**
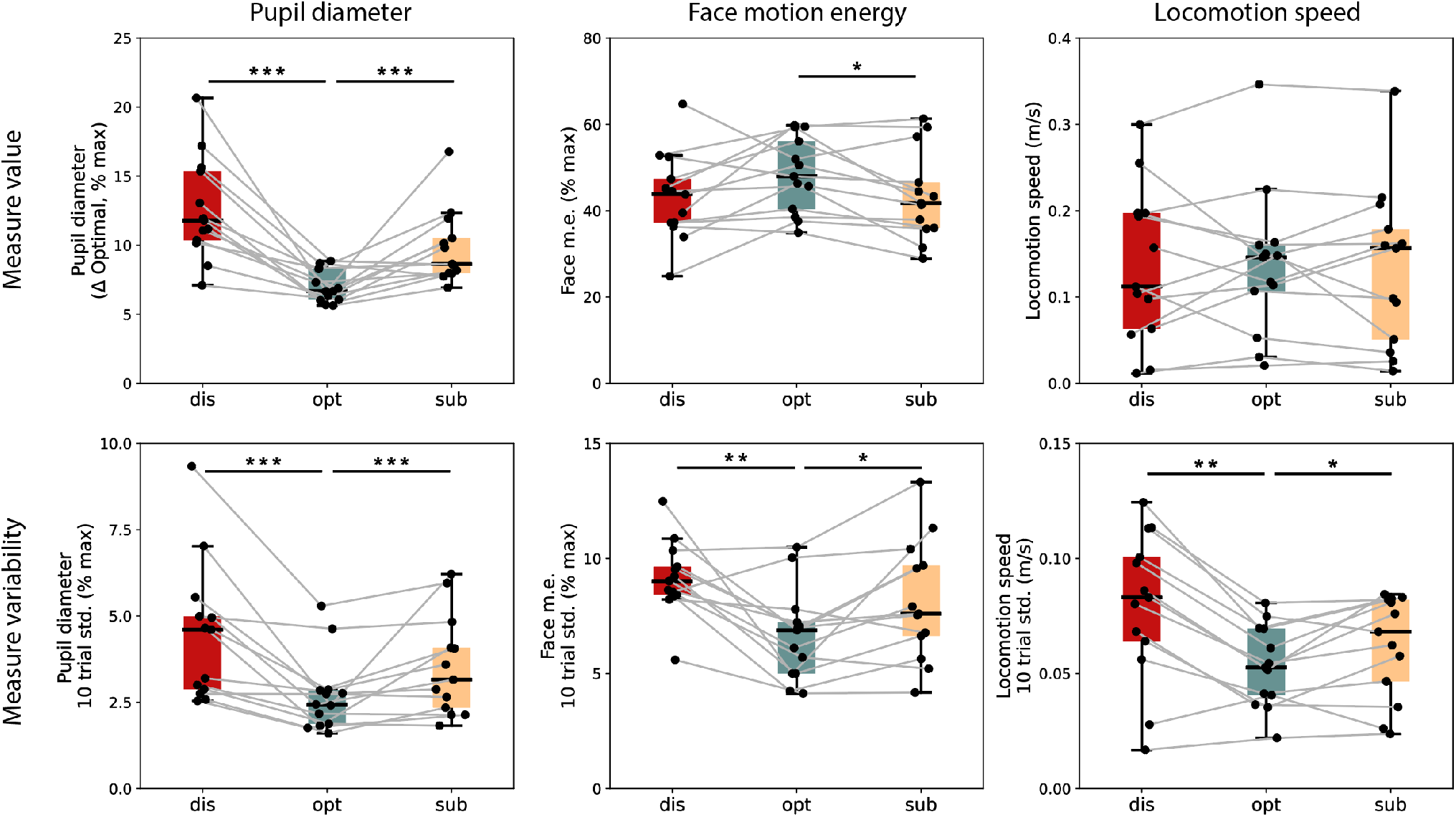
Value and variability of arousal measures by performance states. The average value and standard deviation of the past 10 trials of pupil diameter, face motion energy, and locomotion speed for each subject. Pupil diameter values are the absolute difference from the optimal for each mouse, and are significantly smaller in the optimal state than sub-optimal and disengaged states. Face motion energy is slightly higher in the optimal state, and there is no difference in locomotion speed. The average standard deviation of the past 10 trials is significantly lower during the optimal state for all recorded arousal measures. * indicates p < .05, ** indicates p < 0.01, *** indicates p < 0.001.

**Fig. S5.**
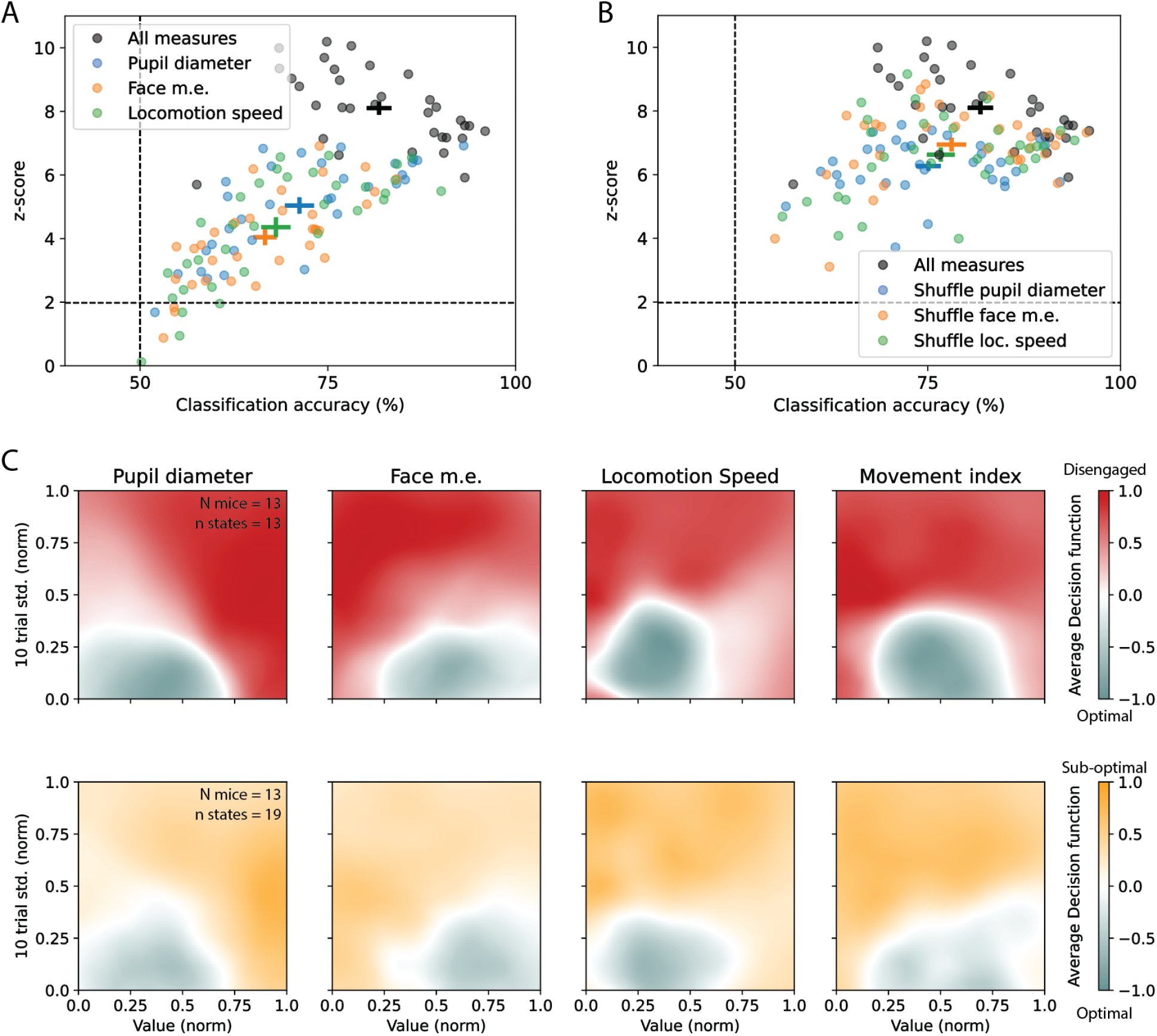
Arousal measure contributions to performance state classification. **(A)** State classification using raw behavioral measures shows that pupil diameter is the best single measure used. **(B)** Shuffling the value and standard deviation of individual arousal measures indicates their unique contributions. Pupil diameter contributes significantly more to decoding than face motion energy (p = 0.004, Wilcoxon signed rank test), but only slightly more than locomotion speed (p = 0.14, Wilcoxon signed rank test). **(C)** Average decision functions across all subjects for disengaged (top) and sub-optimal states (bottom). Note that the low variability consistently predicts optimal state, along with intermediate levels of pupil, locomotion, and overall movement.

